# Oropouche virus infects human neural progenitor cells and alters the growth of brain organoids

**DOI:** 10.1101/2025.10.01.679540

**Authors:** Alexandra Albert, Laurine Couture, François Piumi, Sophie Lebon, Pierre Gressens, Muriel Coulpier, Ali Amara, Vincent El Ghouzzi, Laurent Meertens

**Affiliations:** Université Paris Cité, NeuroDiderot, Inserm UMR 1141, F-75019 Paris, France; Biology and Pathogenesis of Viral Infection Team, INSERM UMR 1342, Institut de Recherche Saint-Louis, Université Paris-Cité, Hôpital Saint-Louis, 75010 Paris, France; UMR Virologie, Anses, INRAE, École Nationale Vétérinaire d’Alfort, Université Paris-Est, Maisons-Alfort, France

## Abstract

Oropouche virus (OROV), is the etiologic agent of Oropouche fever (OROF), an emerging zoonotic disease that has been prevalent in South and Central America since the 1960s. Starting late 2023, the current outbreak has raised concern about vertical transmission and potential adverse pregnancy outcomes. While serological evidences and recent studies support the vertical transmission of OROV, how infection affects the fetal brain remains unclear. Here we show that a strain of OROV, FG_2020, phylogenetically-related to the epidemic strain, efficiently and productively infects human neural progenitor cells and iPSC-derived brain organoids. The main phenotypic effect is proliferation arrest associated with apoptosis leading to the loss of neural rosette organization, a key signature of developing brain architecture. These data indicates that fetal brain is susceptible to OROV infection and that vertical transmission during the first months of pregnancy could lead to pathological effects on brain development.

---

The recent resurgence of Oropouche virus (OROV) raises urgent concerns about its potential to cause congenital disease. OROV is an arthropod-borne virus (arbovirus) from the *Orthobunyavirus* genus transmitted to human primarily by *Culicoides paraensis* midges. First identified in 1961 in Trinidad and Tobago, OROV has remained endemic in the Amazonian Basin^1,2^. However, it has re-emerged in late 2023 causing outbreaks across several countries in South America and the Caribbean, with more than 26,000 confirmed cases as of July 2025^3^.

OROV is a member of the Simbu serogroup, which includes veterinary viruses such as Schmallenberg virus (SBV), Akabane virus and Aino virus, pathogens recognized for their teratogenic effects in ruminants. Mounting evidence now raises concern that OROV may exert similar pathogenic effects during pregnancy in humans. Recent case reports have linked OROV infection to adverse pregnancy outcomes including miscarriages, fetal demise and congenital anomalies such as microcephaly^4–8^. Experimental studies further support this possibility. For instance, OROV has been shown to readily infect human placental explants and replicate efficiently in cytotrophoblasts and syncytiotrophoblasts^9^. Furthermore, in a murine gestational infection model, the virus was shown to cross the maternal-fetal barrier and disseminate in fetal tissues^10^.

Although several case studies have reported neuropathogenic diseases associated with OROV infection^11–14^ and animal model of infection have confirmed the neurotropism of OROV^15–18^, it remains unknown whether OROV can directly infect the developing brain and impair neurodevelopment in human, as documented for Zika virus. Addressing this gap is critical for assessing the potential of OROV to cause congenital disease. Here, we used human neural progenitor cells (hNPCs) and human induced pluripotent stem cell (hiPSC)-derived cerebral organoids as experimental models to define the susceptibility of neural cells to OROV and evaluate the consequences of their infection on early brain development in humans.

To investigate the susceptibility of NPCs to OROV, we inoculated hNPCs derived from first-trimester fetal central nervous system tissue^19^, with increasing multiplicities of infection (MOI) of the FG_2020 strain. This strain was isolated during the 2020 outbreak in French Guiana and clusters with isolates of the current outbreak within the monophyletic clade (OROV_BR-2015-2024_)^20^. As a control we infected under the same condition Vero E6 cells, an IFN-deficient simian cell line known to be permissive to OROV. Infection was quantified 24 hours post-inoculation by flow cytometry using an antibody directed against the viral nucleoprotein protein (N). In both hNPCs and Vero E6 cells, the proportion of infected cells increased in an MOI-dependent manner (Figure 1A). Notably, hNPCs displayed a higher infection rate than Vero E6 cells, demonstrating their marked susceptibility to OROV. Infection of hNPCs was also observed following inoculation with the prototypical BeAn19991 strain, isolated in 1960 (Figure S1A), albeit at lower levels, ruling out the possibility that neurotropism is a recently acquired feature of the emerging strain. Furthermore, productive replication in hNPCs was detected as early as 8 hours post-infection (hpi) with the release of infectious viral particles in the supernatant, which reach a plateau at 24 hpi (Figure 1B). By 48 hpi, OROV induced a pronounced cytopathic effect relative to mock-infected cells (Figure 1C and Figure S1B).

**Figure 1.**
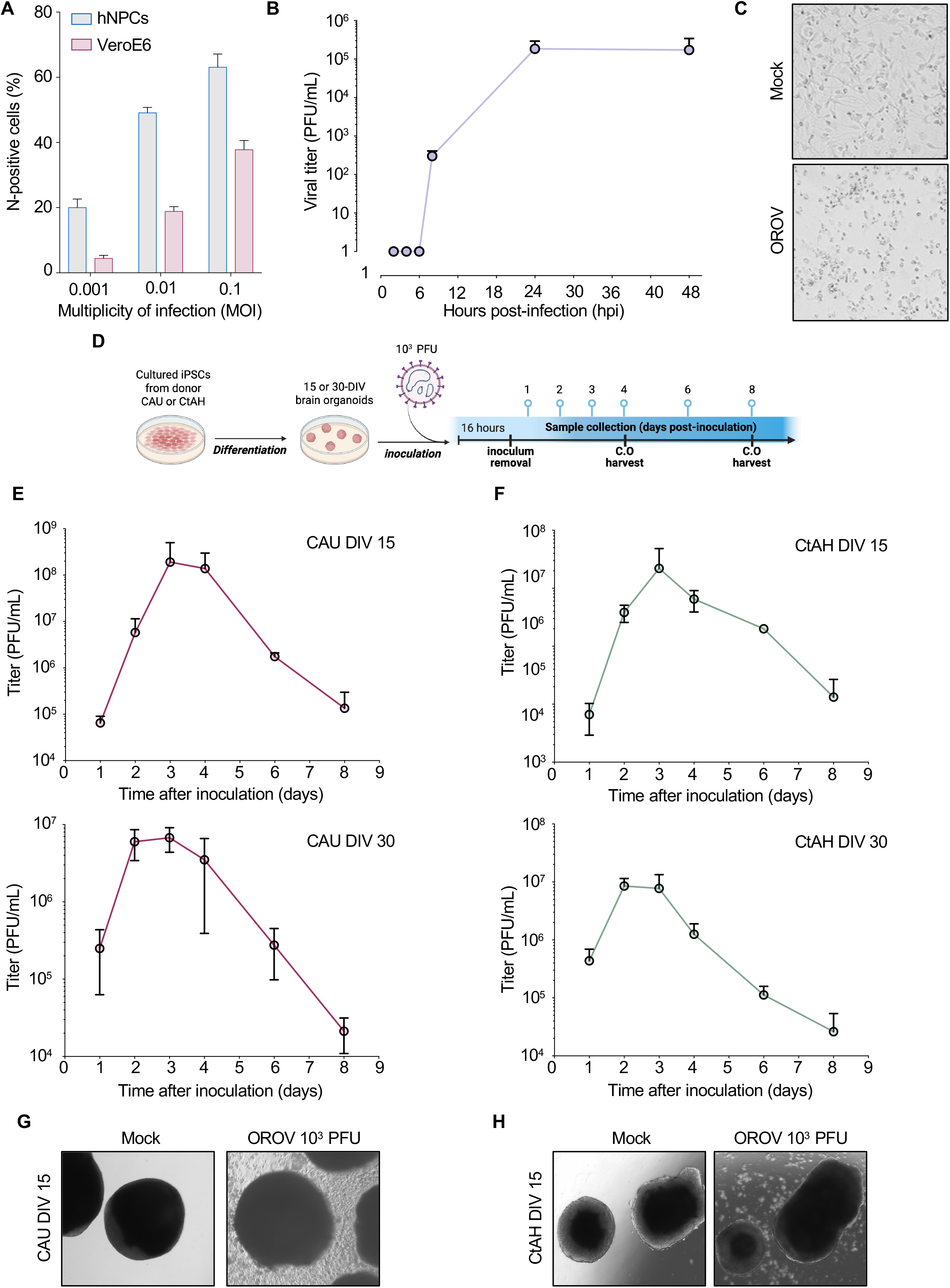
OROV productively infects human NPCs and forebrain organoids. (A) hNPCs and VeroE6 cells were inoculated with increasing MOI of OROV_FG_2020. Percentage of viral antigen-positive cells was assessed 24 hpi by flow cytometry. Data shown are mean ± SD of three independent experiments in duplicate. (B) hNPCs were inoculated with OROV (MOI 0.01) and inoculum was replaced with culture medium 2 hours post-infection. Supernatant were collected at different time points and titrated by plaque assay on VeroE6 cells. Data shown are mean ± SD of two independent experiments in duplicate. (C) Brightfield images from hNPCs infected with OROV (MOI 0.01) for 48 hours. Data shown are representative of three independent experiments. (D) Schematic of iPSC-derived forebrain-organoids infection with OROV. Inoculum was replaced 16 hpi with culture medium and supernatant was collected at indicated time points. (E and F) Multistep growth curves of OROV in iPSC-derived forebrain-organoids from donors CAU (E) and CtAH (F). Progeny virus released in supernatants collected at different time points were titrated by plaque assay on VeroE6 cells. Data shown are mean ± SD; n=2 independent experiments in duplicate. (G and H) Representative brightfield images of mock and OROV infected DIV-15 forebrain organoids at 4 dpi.

We next evaluated whether OROV could establish productive infection in early-stage brain organoids (15 or 30 days in vitro, DIV), where the majority of neuroepithelial progenitors express SOX2 and actively divide around the ventricle, forming the ventricular zone (VZ). To induce robust neural differentiation of hiPSCs, we cultured the cells in suspension in the presence of dorsomorphin and SB-431542, two inhibitors of the SMAD pathway that facilitate neuralization and produce a default dorsal forebrain fate as previously described^21^. Forebrain organoids derived from two independent iPSC lines (CAU and CtAH) were inoculated with 10^3^ PFU of OROV (Figure 1D). Viral progeny in culture supernatants was quantified by plaque assay at multiple time points post-infection. For both donor lines, infectious titers were detectable at 24 hpi, peaking at 2 dpi in DIV-30 organoids and 3 dpi in DIV-15 organoids (Figure 1E–F), before declining sharply by 4 dpi. At this time point, morphological assessment revealed abundant cellular debris in OROV-infected cultures compared to mock controls (Figure 1G–H), consistent with virus-induced cytopathology. Together, these results demonstrate that OROV productively infects neural progenitors and induces cytopathic effects in developing human brain organoids.

To assess the impact of OROV infection on the architectural organization of brain organoids, we performed immunofluorescence analysis on sections stained for the neural progenitor marker SOX2 and OROV-N antigen. In mock-infected controls, early-stage organoids displayed the characteristic SOX2+ neural progenitor organization in neural rosette (Figure 2A). By 4 dpi, OROV antigen was readily detected within SOX2+ progenitors (Figure 2A), coinciding with neural rosette disruption. At 8 dpi, OROV antigen was widespread accompanied by a complete loss of organized brain-like structures, in contrast to the preserved architecture in mock-infected organoids. These findings indicate that OROV targets neural progenitors and disrupts structural organization during early brain development.

**Figure 2.**
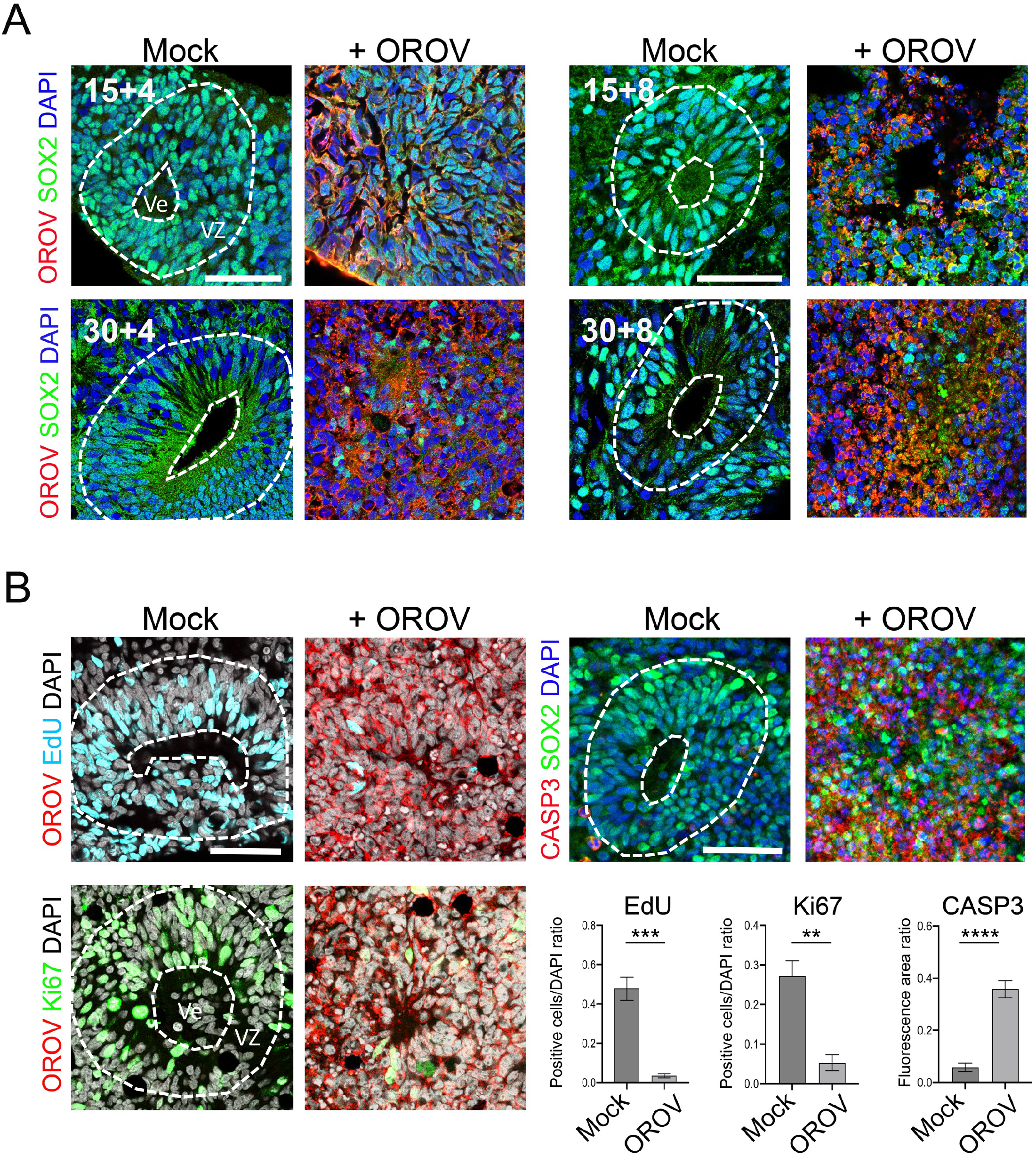
Characterization of the expression of neural, proliferation and cell death markers in OROV-infected early forebrain organoids. (A) Immunofluorescent staining representative of the ventricular zone (VZ) of mock or infected-organoid showing progenitors at DIV-15 and DIV-30 labelled with SOX2 (green) surrounding the ventricle (Ve), and the presence of OROV (red) 4- or 8-days post-infection. Scale Bar 50 μm. (B). Immunofluorescent staining and representative quantification of the ventricular zone (VZ) of mock or infected-organoid at DIV-30 showing the proportion of cells that incorporated EdU for 3 hours prior to fixation (in cyan), cells in cycle (Ki67+ in green), and the cleaved caspase 3 fluorescent signal per unit area (in red, right panel). Data are presented as mean ± SEM. *p≤0.05 (Student’s t-test). Images shown are representative of two independent experiments. Scale Bar, 50 μm.

We further assessed the impact of infection on progenitor proliferation by quantifying incorporation of the thymidine analog EdU, incubated for 3 hours prior to DIV-30 organoid fixation. At 4 dpi, OROV-infected organoids showed a 13.7-fold reduction in EdU+ cells per section compared to mock controls (Figure 2B). Consistent with this observation, the proportion of cells positive for the proliferation marker Ki67 was reduced approximately five-fold (Figure 2B).

To determine whether OROV infection could induce neural progenitor cell death via apoptosis, DIV-30 organoid sections were stained for activated caspase-3 (CASP3). While mock-infected organoids showed minimal CASP3 signal, OROV-infected organoids exhibited a marked increase in CASP3^+^ cells (Figure 2B), indicating virus-induced apoptosis. Combined, these results provide strong evidence that OROV infection impairs neural progenitors proliferation and survival, thereby compromising early stages of human brain development.

Our study demonstrates that Oropouche virus (OROV) productively infects human neural progenitor cells (hNPCs) in 2D-culture and 3D-forebrain organoids, leading to profound cytopathic effects characterized by reduced proliferation, apoptosis and loss of structural organization. At the very beginning of corticogenesis, these progenitors are highly polarized neuroepithelial cells that actively self-amplify and play a decisive role in the neural fate of the radial glial cells that derive from them and their descendants, which give rise to both neurons and glia^22^. While OROV infection was previously known to infect neurons or microglial cells in adult brain slices^23^, our findings identify early neural progenitors as a primary cellular target of OROV and suggest that infection during early neurodevelopment may severely compromise brain development. The ability of OROV to infect fetal-derived progenitors and cerebral organoids strengthens the hypothesis that OROV may induce congenital disease, raising significant public health concerns given its endemic and epidemic activity in South America.

The neuropathogenic features we describe for OROV bear striking similarities to those reported for Zika virus (ZIKV)^24–26^. ZIKV is firmly established as a teratogenic agent responsible for congenital Zika syndrome (CZS)^27^, which has also been directly linked to the occurrence of congenital microcephaly, with a particular tropism for NPCs located in the ventricular zone (VZ) and the sub-ventricular zones^26^. In these cells, ZIKV infection induces capase-3 activation, cell death, and reduced proliferation, leading to cortical thinning and impaired brain growth^26,28,29^. Similarly, we demonstrate that OROV infects SOX2^+^ progenitor populations, disrupts rosette-like structures that mimic early VZ, and induces apoptosis alongside impaired proliferation. These phenotypes are hallmarks of impaired corticogenesis and provide a plausible mechanistic basis for OROV-induced neurodevelopmental defects, including congenital microcephaly. Comparative analysis of OROV and ZIKV are therefore likely to reveal both shared pathways and virus-specific mechanisms of virus-induced teratogenesis that could be targeted for therapeutical intervention or public health prevention strategies.

Mechanistically, we observed an early peak in viral titers in organoid supernatants followed by a decline coinciding with widespread cell death. This suggests a model in which OROV rapidly depletes progenitor pools via apoptosis and growth arrest, and where subsequent collapse of brain architecture irreversibly impairs neurogenesis. This pattern resembles the “hit-and-run” model of ZIKV neuropathogenesis, in which transient replication leads to long-lasting injury to progenitor populations. However, alternative explanations remain possible: declining titers may directly reflect necrosis of brain tissue due to intensive viral dissemination and apoptosis, potentially associated with more severe outcomes such as miscarriage or fetal demise. These findings raise the possibility that maternal and fetal host factors restricting viral spread may critically influence the severity of vertical transmission outcomes.

A key determinant of whether OROV infection of fetal brain occurs *in vivo* is the ability of the virus to cross the placental barrier. Vertical transmission requires viral entry into placental cells, evasion of innate immune responses at the maternal-fetal interface, and subsequent access to the fetal compartment. ZIKV studies have shown that trophoblasts, Hofbauer cells, and placental endothelial cells are all susceptible, facilitating viral passage to the fetus^30–33^. Recent experimental work using human placental explants and trophoblast models has reported OROV replication in placental tissues, supporting a similar mechanism of vertical transmission^9^. Animal studies have further suggested preferential infection of first-trimester placentas^10^. This is consistent with the association between the higher risk of congenital diseases when ZIKV infection occurred at the end of the first trimester of pregnancy. Both viruses may exploit the permissive environment of this developmental stage, when the placental barrier is still maturing and neural progenitors are undergoing rapid proliferation. Future development of *in vivo* pregnancy models that recapitulate human maternal–fetal immunobiology will be essential to delineate the timing, efficiency, and outcomes of OROV vertical transmission and to define its potential role in congenital Oropouche syndrome.

Finally, our findings align with epidemiological observations and comparative virology within the Simbu serogroup. Related orthobunyaviruses, such as Akabane and Schmallenberg viruses, cause severe fetal anomalies in ruminants, including arthrogryposis, cerebellar hypoplasia, and hydrocephalus^4^. These similarities suggest that OROV may share conserved mechanisms of teratogenesis with these animal pathogens. While definitive clinical evidence of congenital Oropouche syndrome in humans is still lacking, the strong similarities with CZS highlight the urgent need for enhanced surveillance, targeted experimental studies, and preparedness in regions where OROV is endemic or emerging.

## Supporting information

Supplementary Informations

## Acknowledgements

We would like to thank Drs. Pierre Vanderhaeghen and Odile Blanchet for providing us with the CtAH line and the hNPC, respectively, as well as the Paris Brain Child Institute (ICE) for its support of NeuroDiderot’s Human Brain Organoid core facility (HumBO). This work has received support under the program “Investissement d’Avenir” launched by the French Government and implemented by ANR, under the reference « ANR^−^18^−^IdEx−0001 » as part of its program « Emergence », and under the reference ANR-23-IAHU-0010 as part of the France 2030 program. L.M is supported by the Laboratoire d’Excellence “Integrative Biology of Emerging Infectious Diseases” (grant n°ANR-10-LABEX-62-IBEID).

