## Supplementary Informations for "Oropouche virus infects human neural progenitor cells and alters the growth of brain organoids"

### Supplemental Figures

**A**

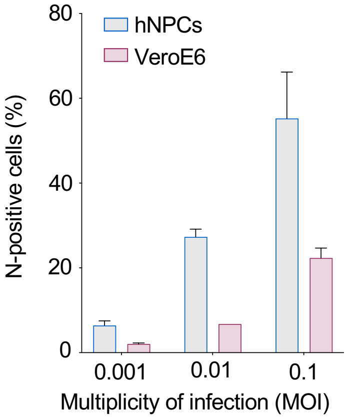


**B**

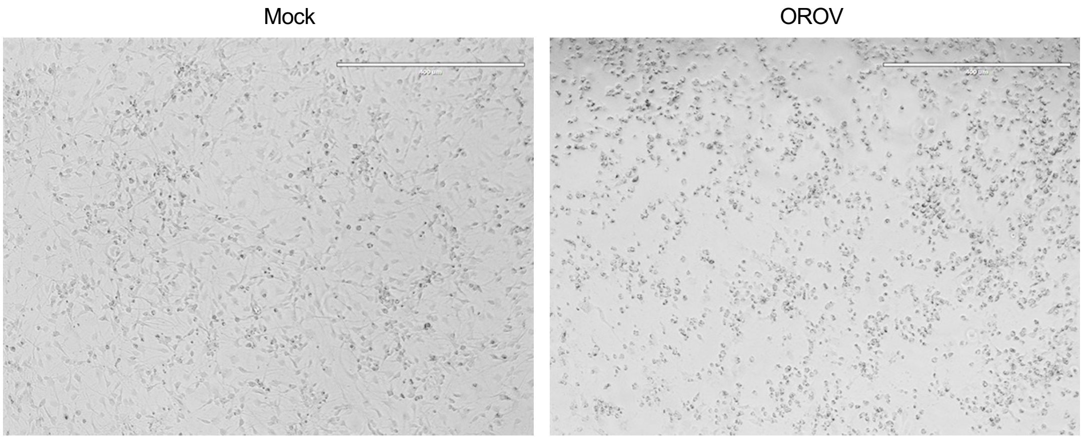


**Figure S1. hNPCs are susceptible to infection with prototypical strain BeAn1991, related to Figure 1.** (A) hNPCs and VeroE6 cells were inoculated with increasing MOI of OROV_BeAn19991. Percentage of viral antigen-positive cells was assessed 24 hpi by flow cytometry. Data shown are mean ± SD of three independent experiments in duplicate. (B) Brightfield images from hNPCs infected with OROV (MOI 0.01) for 48 hours. Data shown are representative of three independent experiments. Scale bar 400 μm.

### Experimental Procedures

#### Cell culture

Vero E6 (gift of Marc Lecuit, Pasteur Institute Paris, France) cells were grown in DMEM supplemented with 10% FBS, 1% P/S, 1X Glutamax and 25 mM HEPES. Hamster BSR-T7/5 cells (gift of Alain Kohl, MRC-University of Glasgow, Scotland), which stably express the T7 RNA polymerase, were grown in GMEM supplemented with 10% (FBS), 1% P/S, 10% tryptose phosphate broth (Thermo Fisher Scientific), 1% GlutaMAX and Geniticin (1mg/ml). Human neural progenitor cells (hNPCs) were prepared and cultured as previously described[19] .All cell lines were cultured at 37°C in presence of 5% $CO_2$. All cells were routinely tested to be free of mycoplasma contamination.

#### Ethics approval and consent to participate

Human fetus was obtained after legal abortion with written informed consent from the patient. The procedure for the procurement and use of human fetal central nervous system tissue was approved and monitored by the “Comité Consultatif de Protection des Personnes dans la Recherche Biomédicale” of Henri Mondor Hospital, France. All methods were in compliance with relevant French laws and institutional guidelines. Authorization and declaration numbers from the French Research Ministry are AC-2017-2993 (CHU Angers) and DC-2019-3771 (UMR Virologie, Maisons-Alfort).

#### Culture of hiPS cell lines and forebrain organoid differentiation.

The hiPSC lines, PCi-CAU1 (male donor) and CtAH (female donor), were used in this study. PCi-CAU1 cells were purchased from Phenocell (Grasse, France) and CtAH were generously provided by P. Vanderhaeghen’s lab. Cells were maintained in 6-well plates in feeder-free conditions on Matrigel (BD/Corning) using StemMACS$^{TM}$ iPS-Brew XF (Miltenyi Biotec) and non-enzymatically dissociated with StemMACS Passaging Solution XF (Miltenyi Biotec). All cells were cultured at 37 °C under 20% O2 and 5% CO2. Forebrain-specific organoids were generated using established protocols as previously described[21]. Briefly, human iPSC colonies were detached from the well with 0.35 mg/ml Dispase (Invitrogen) treatment for 35 min and suspended in neural induction medium, consisting of DMEM/F12 (Life Technologies), 20% KO serum replacement (Life Technologies), 1% MEM-NEAA (Life Technologies), 1% Glutamax (Life Technologies), 1% Penicillin-Streptomycin (Life Technologies), 2-mercaptoethanol (Sigma), supplemented with 5 µM Dorsomorphin (Sigma) and 10 µM SB-431542 (Tocris) in ultra-low attachment plates for 6 days under agitation with a daily medium change. On day 6, neural spheroids were grown in neural differentiation medium consisting of Neurobasal A (Life Technologies), 2% B27 (Life Technologies), 1% Glutamax (Life Technologies), 1% Penicillin-Streptomycin (Life Technologies) supplemented with 20 ng/mL EGF (R&D Systems) and 20 ng/mL FGF-2 (R&D systems), with daily medium change until day 15, then every other day. From day 25 onwards, EGF and FGF-2 were replaced by 20 ng/mL BDNF (PeproTech) and 20 ng/mL NT3 (PeproTech) and medium was changed every two-three days.

**Virus strains and culture**

OROV_FG_2020 (strain French Guiana; provided by D. Missé, IRD, Montpellier) was propagated with limited passage on Vero E6 cells. Plasmids pTVTOROVL, pTVTOROVM and pTVTOROVS that encodes the antigenomic sense cDNA of the corresponding viral genome segments used for OROV rescue, were a kind gift of Benjamin Brennan (MRC-University of Glasgow, Scotland). The cDNAs clones were constructed from the OROV strain BeAn19991, isolated in 1960 from a sloth (*Bradypus tridactylus*) in Brazil. To recover infectious virus OROV from the cDNAs, BSR-T7/5 (2 x $10^6$ cells per 60 mm culture dishes) cells were transfected with 2 $\mu$g of the pTVTOROVL, pTVTOROVM, and pTVTOROVS plasmids with 13 $\mu$L of lipofectamine 3000 following the manufacturer's recommendations. Supernatant was harvested 5 days after transfection, clarified by centrifugation and used for viral propagation on Vero E6 cells to generate the viral stocks.

For all the viral stocks, viruses were purified through a 20% sucrose cushion by ultracentrifugation at 80,000*g* for 2 hours at 4 °C. Pellets were resuspended in HNE1X pH 7.4 (HEPES 5 mM, NaCl 150 mM, EDTA 0.1 mM), aliquoted and stored at −80 °C. Viral stock were titrated on Vero E6 cells by plaque-forming assay and virus titers are expressed as plaque-forming units (PFU) per mL.

**Infection assay**

For infection quantification by flow cytometry analysis, hNPCs were plated in 24-well plates (1.6 x $10^5$ cells/well) and Vero E6 were plated in 12-well plates (8 x $10^4$ cells/well). Cells were inoculated for 24 h, trypsinized and fixed with 2% (v/v) paraformaldehyde (PFA) diluted in PBS for 15 min at room temperature. Cells were incubated for 1 hour at 4 °C with anti-N rabbit purified antibody for OROV. Antibodies were diluted in permeabilization flow cytometry buffer (PBS supplemented with 5% FBS, 0.5% (w/v) saponin, 0.1% sodium azide). After washing, cells were incubated with 1 µg $mL^{-1}$ of Alexa Fluor 647-conjugated secondary antibody diluted in permeabilization flow cytometry buffer for 30 min at 4 °C. Acquisition was performed on an Attune NxT Flow Cytometer (Thermo Fisher Scientific) and analysis was done by using FlowJo software (TreeStar).

To quantify infectious viral particles released during hNPCs infection, cells were inoculated for 2 hours with viruses, washed once and then maintained in culture medium over a 48-hour period. At indicated time points supernatants were collected and kept at -80°C. Virus titers were determined on Vero E6 cells by plaque-forming assay and expressed as plaque-forming units (PFU) per mL.

To quantify infectious viral particles released during hiPS-derived forebrain organoid infection, 15- or 30-DIV organoids were divided evenly in low attachment p24-well plates (4-5 organoids per well). Organoids were inoculated 16 hours with $10^3$ PFU/well under continuous agitation (70 rpm), washed twice and then maintained in culture medium over an 8-day period under agitation. Culture medium was replaced every 24 hours. Starting at 8 hours after media replacement (1 dpi), supernatants were collected every 24 hours and kept at -80°C. Virus titers were determined on Vero E6 cells by plaque-forming assay and expressed as plaque-forming units (PFU) per mL. At 4 and 8 dpi, organoids were fixed in 4 % paraformaldehyde for 20 min at 4°C and cryoprotected in 30 % sucrose in PBS.

**Titration assay**

Vero E6 cells were plated on 24 well-plates (2 x $10^5$ cells/well) and incubated with 200 µL of 10-fold serial dilution of supernatant collected from OROV infected cells. After 40 min, the inoculum was replaced with Avicel 2.4% mixed at equal volume with DMEM supplemented with 4% FBS, 2% Glutamax, 50mM $MgCl_2$, 0.225 % of $NaHCO_3$, and incubated 3 days at 37°C. Then, Vero E6 cells were washed twice with phosphate-buffered saline (PBS) and stained with an aqueous solution containing 1% formaldehyde, 0,5% crystal violet, and 0,89% sodium chloride, for 15 min at room temperature. Vero E6 cells were then washed with water to visualize lysis plaques.

**Immunofluorescent staining.**

Three hours prior to fixation, the organoids were incubated in the presence of 5-ethynyl-2'-deoxyuridine (EdU) according to the kit manufacturer's instructions (Pharmingen™ 647 EdU Click Proliferation Kit). After fixation in 4% PFA, the organoids were immersed in a 30% sucrose PBS solution, then embedded in 7.5% gelatin/15% sucrose blocks and cryopreserved before being cryo-sectioned into 20 µm serial sections. For immunocytochemical staining, the sections were first immersed in a PBS solution containing 0.1% Tween, incubated for 60 minutes in a blocking solution containing 0.3% Triton, 6% donkey serum in PBS, and then overnight at 4°C in a solution containing 0.1% Triton, 6% donkey serum in PBS and primary antibodies directed against SOX2, OROV anti-N, Ki67 or Cleaved Caspase-3. After three washes in PBS, the samples were incubated for 60 minutes at room temperature with fluorescent secondary antibodies at a dilution of 1:500 (Jackson Immunoresearch) in the same solution as the previous day, rinsed, incubated in DAPI and mounted with DAKO fluorescence mounting medium. For EdU labelling, the sections were fixed again in 4% PFA right after immunocytochemistry, rinsed and then incubated for 30 min at room temperature in the reagent containing the catalyst, reaction buffer and far-red dye according to the kit manufacturer's instructions, rinsed again and then counterstained with Dapi and mounted with Dako fluorescence medium.

**Image analysis and statistics**

Immunofluorescent sections were analysed using a Leica TCS SP8 confocal scanning system (Leica Microsystems). Eight-bit digital images were collected from a single optical plane using a 20x HC PL APO CS2 oil-immersion Leica objective (numerical aperture 0.75) or a 40x HC PL APO CS2 oil-immersion Leica objective (numerical aperture 1.30). For each optical section, double- or triple-fluorescence images were acquired in sequential mode to avoid potential contamination by linkage specific fluorescence emission cross-talk. Settings for laser intensity, beam expander, pinhole (1 Airy unit), range property of emission window, electronic zoom, gain and offset of photomultiplicator, field format, scanning speed were optimized initially and held constant throughout the study so that all sections were digitized under the same conditions. All images were analysed with ImageJ and statistical analysis was performed using GraphPad Prism 8.0 (GraphPad Software, San Diego, CA, USA). Mean values for the experimental groups were calculated and statistically analysed by two-tailed unpaired Student's *t*-test for single comparisons to control. The values of $P < 0.05$ was considered statistically significant. Data are presented as mean ± SEM.
